# TEPEAK : A novel method for identifying and characterizing polymorphic transposable elements in non-model species populations

**DOI:** 10.1101/2023.10.13.562297

**Authors:** Devin Burke, Edward Chuong, William Taylor, Ryan Layer

## Abstract

Transposable elements (TEs) replicate within genomes and are an active source of genetic variability in many species. Their role in immunity and domestication underscores their biological significance. However, analyzing TEs, especially within lesser-studied and wild populations, poses considerable challenges. To address this, we introduce TEPEAK, a simple and efficient approach to identify and characterize TEs in populations without any prior sequence or loci information. In addition to processing user-submitted genomes, TEPEAK integrates with the National Center for Biotechnology Information (NCBI) Sequence Read Archive (SRA) to increase cohort sizes or incorporate proximate species. Our application of TEPEAK to 256 horse genomes spanning 11 groups reaffirmed established genetic histories and highlighted disruptions in crucial genes. Some identified TEs were also detectable in species closely related to horses. TEPEAK paves the way for comprehensive genetic variation analysis in traditionally understudied populations by simplifying TE studies. TEPEAK is open-source and freely available at https://github.com/mrburke00/TEPEAK.

## Introduction

While there is a growing appreciation for the impact structural variation (SV) has on genetic and phenotypic diversity, accurately identifying the many types of structural variation remains challenging. Among SVs, transposable elements (TEs) are no exception. TEs are mobile genetic elements which can independently insert into alternate genomic regions thereby multiplying themselves throughout the host genome, occupying a substantial portion of many species’ genomes (Wells 2020). Despite their significance, and the availability of a large number of sequenced samples, TEs are poorly cataloged in many non-model species, including those that remain a source of SVs. This gap is due in part to the lack of simple and efficient methods available to identify and accurately differentiate polymorphic and novel TE insertions.

TEs are observed in nearly every eukaryotic genome and have profound implications due to the unique biological characteristics and the complementary relationship between environmental and population genetic factors. For example, insertions of TE sequences into coding regions can cause expansion and propagation of novel genes (Baduel 2021, Kidwell 1997) and drive adaptive phenotypic variation from environmental pressures like global warming, disease and the adaptive immune system, speciation, and inbreeding and endangered populations (Baduel 2021, Platt 2016, De Kort 2022). Many types of TEs demonstrate locational specificity, with some targeting regions that minimize deleterious effects and others maximizing their propagation (Baduel 2021, Richardson 2015). These behaviors are likely due to the evolutionary stability of TEs (Bourgeois 2019). While TE sequences decay over time, often resulting in the loss of functionality, they often retain promoters and other remnant genomic features that indirectly impact nearby genes (Richardson 2015), including rewiring entire gene networks (Kidwell 1997).

TEs exhibit remarkable diversity, with over 273,000 different families in the most recent DFAM (3.3) database (Storer 2021). These families are distinguished into two different major classes; class I - retrotransposons and class II – DNA transposons. Class I elements involve an RNA intermediate, which undergoes reverse transcription yielding a DNA copy that is inserted into a new location (Kidwell 1997, Richardson 2015, Bourque 2018). Class II elements move via a “cut-and-paste” mechanism wherein a transposase enzyme directly mediates the joining of the DNA intermediate to its new site (Batcher 2023, Bourque 2018). These two major classes are further divided into superfamilies, families, and subfamilies/subclasses (Kidwell 1997, Richardson 2015).

Traditionally, TE families are defined as sequences that share 80% coverage with 80% identity and are thus most often characterized by phylogenetic relationships (although there are families where such characterization is not adhered to) (Storer 2021, Wells 2020). Subclasses are then classified by mechanisms of replication or integration. They can be broadly distinguished by whether the element encodes for enzymes which facilitate self-transposition (Kidwell 1997, Wells 2020, Bourque 2018). However this classification does not encapsulate TE families that have emerged ‘de novo’ as is the case with short interspersed nuclear elements (SINEs) that are often derived from noncoding RNAs. Many of these SINE elements in fact hijack long interspersed nuclear elements (LINEs), which do encode for autonomous transposition (Richardson 2015). It has been shown that SINEs often share homology with LINEs, thereby explaining the capacity for LINEs to reverse transcribe and integrate SINEs (Richardson 2015).

TE discovery can roughly be divided into identification of polymorphic insertions of previously annotated TE families and insertion of novel unannotated TE families (Platt 2016). Current polymorphic insertion methods use both sequence homology-based and ‘de-novo’ structural feature searches to identify polymorphic insertions from a sequence or structural motif of interest (Platt 2016, Abrusan 2009, Feschotte 2009, Gardner 2017, Smit 2010, Xu 2007). However, these methods struggle to identify TEs different from those previously identified (Hoen 2015). Furthermore, all TEs, to varying degrees, suffer from decay over time which complicates the ability to extract all polymorphic insertions of the same family (Wells 2020). Using expert domain knowledge or ‘de novo’ methods is usually required to identify novel unannotated TE families (Hoen 2015). This bottleneck results in a small and isolated perspective of the TE landscape in a species (Domínguez 2020).

Another consequence of TE sequence decay is the prevention of TE annotation processes from entirely avoiding manual curation (Platt 2016). Both homology-based and ‘de-novo’ based structural feature searches struggle to identify TEs distinct from those already well established and described (Storer 2021, Bourgeois 2019). Current methods also often require expert knowledge of the TE landscape in the species of interest (Hoen 2015, Charlesworth 1997, Kidwell 2002, Chen 2020). Researchers can utilize a sequence or transposon structural motif to extract polymorphic TE insertions in samples, but this yields only a small isolated perspective of the TE landscape and often fails to discover heavily mutated sequences and novel TEs altogether (Platt 2016, Hoen 2015, Storer 2021).

Extracting polymorphic insertions remains a critical component of constructing the entire TE landscape for a species. However, the bottleneck of current methods lies in their resulting narrow perspective. Most current methods require the user to provide a list of one or several TE family consensus sequences to detect polymorphic insertions from that family (refs–MELT, TLDR, GRAFFITE, etc). Focusing on only known TE targets in a single species prevents a comprehensive and unbiased analysis of TE evolutionary relationships (Storer 2021). In addition to the high frequency of TE involved horizontal transfers, TE families have a complex evolutionary history. This means that single species studies are often insufficient to fully understand a specific TE’s significance (Domínguez 2020). Without comparing closely related species, information pertinent to a TE’s evolution, frequency, impact, and even fixation in a species cannot be obtained (Serrato-Capuchina 2018, Schrader 2014). Recent advances have allowed for a drastic increase in the rate of new reference assembly releases, which has created a dilemma as the number of releases far outpaces annotation efforts (Storer 2021). This means that TE landscapes of non-model organisms are poorly characterized.

Even with the emergence of more TE focused population genetic studies, the lack of a streamlined process to effectively compare TEs in a population makes it difficult to generalize findings or even carry perspectives learned to other studies (Mérel 2020, Errbii 2021, Domíguez 2020, Zhao 2023, Mezzasalma 2023, Ren 2022, Caballero-López 2022, Shao 2019). There is a need for a method that yields the entire landscape of polymorphic TEs in a species. Such a method would have the potential to not only discover novel TEs, but also highly polymorphic TEs that may not be apparent with the limited focus of other methods. This method should also seamlessly allow comparison with nearby species to augment a near complete understanding of the significance of all TEs in the landscape. Together this method would allow for phylogenetic analysis of multiple TEs across multiple species.

We present TEPEAK, a simple streamlined process that is able to identify polymorphic insertions of annotated TEs and novel TEs. The process is based on detecting overrepresentation of SVs of specific lengths, which are likely to correspond to polymorphisms derived from TEs. TEPEAK uses a peak finding algorithm to identify clusters of TE families showing evidence of polymorphism from structural variant insertion data without excluding low copy number TEs. We then use these clusters of TE families to identify all relevant loci in the population. The output of our process is a simple file containing shared loci among the population as well as polymorphic sequences between samples at the same loci. These clusters of TE families can then be used to compare with any other species inputted into the pipeline.

We demonstrate TEPEAK’s utility by reporting on the TE landscape of 265 horse genomes. We also show that our method identifies polymorphic insertions of the same Equid TE families in black rhino, wild ass, and plains zebra, several of which have not been previously annotated. Finally, we demonstrate our method is able to identify novel TEs in non-model species like black rhino.

## Methods

TEPEAK (Figure 1) begins with the user’s choice of considering automated or customized input. The automated input requires a species upon which the National Center for Biotechnology Information (NCBI) Sequence Read Archive (SRA) is queried for high quality reads and aligned to the respective reference genome (Sayers 2022). The customized input requires prepared BAMs/CRAMs. After insertion calling on each sample, the peak finding algorithm identifies significant peaks in a population wide insertion size frequency plot and automatically queries the consensus sequence for insertions that match the peak size against the DFAM database (Storer 2021). Users can also query custom ranges from the insertion size frequency plot. Once a range of insertion sizes is selected the workflow proceeds to extract all loci in the population with insertion sizes that intersect with the given range. This collection of sequences then undergoes clustering to group highly similar sequences. From here users can perform gene annotations, functional enrichment and sequence phylogenetic analysis. The final output is a pair of files that contain individual loci and merged population loci information.

**Figure 1:**
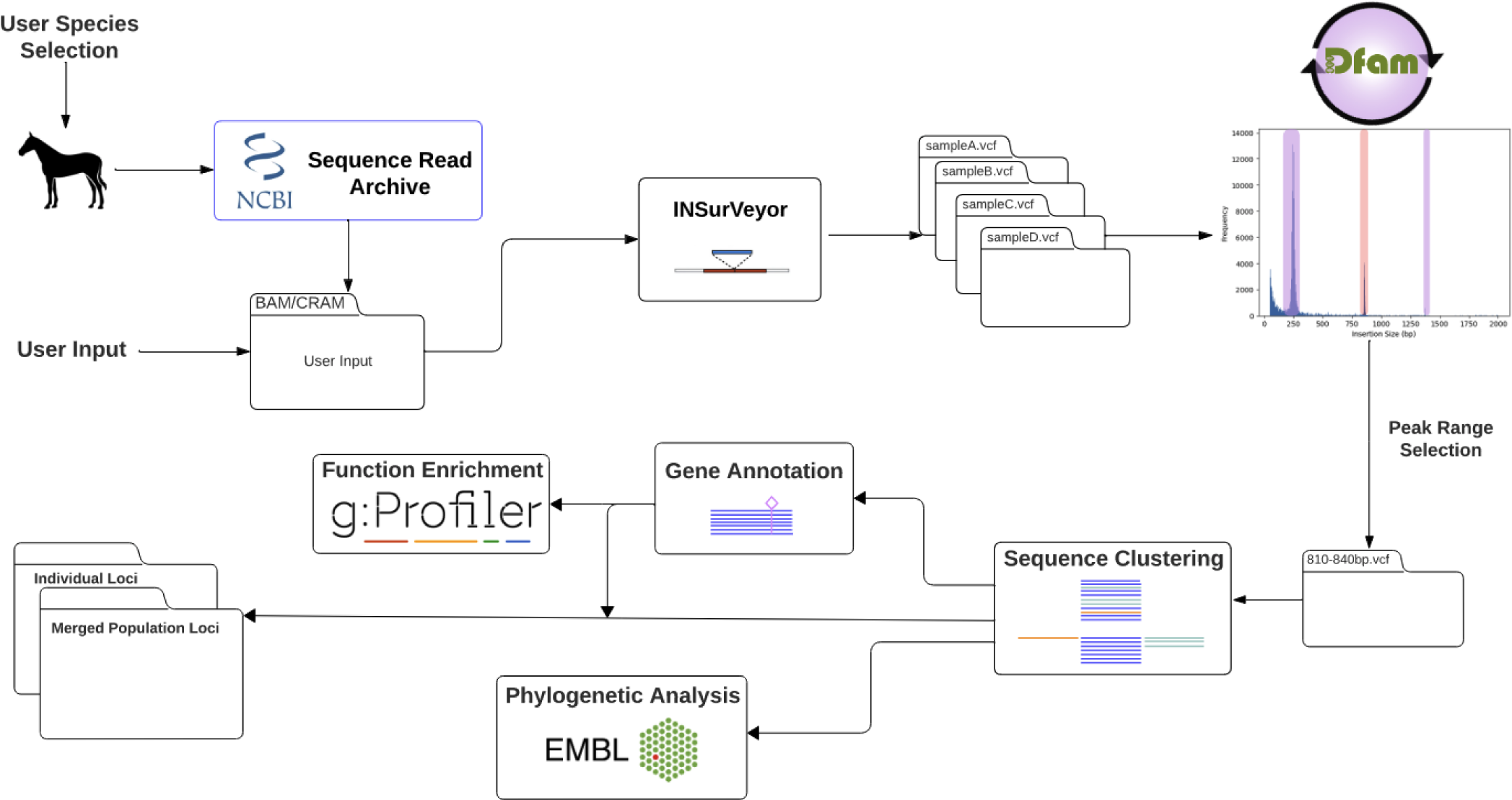
The TEPEAK workflow.

### Data Collection and Preprocessing

The workflow begins by taking user-provided inputs in the form of a species, list of SRA accessions, or pre-aligned Binary Alignment/Map (BAM) files for Illumina short-read sequencing data. If species names are provided, the (NCBI) SDK is utilized to extract a user-defined number of the highest quality SRA accessions available. If SRA accessions are provided, the corresponding raw sequencing data is retrieved and aligned to the user provided reference assembly using BWA Mem (Li 2009). In the case of BAM files, these are directly used as inputs for InSurveyor (Rajaby 2023). In all cases it is recommended that the reference assembly be chromosome level representation and proper quality control measures are utilized on the samples using a suitable alignment tool.

### InSurVeyor (Insertion Calling)

INSurVeyor is a state of the art insertion structural variant calling tool for Illumina short-read sequencing data with sensitivity benchmarks on par with long reads callers (Rajaby 2023). The outcome of this stage is a putative set of TE insertion or SV INDEL sites and sequences for each sample in a Variant Call Format (VCF) file and a summary file detailing the number of insertions per sample. This summary file can be used to further filter poor quality samples.

### Peak Finding Algorithm

The resulting VCF files for each sample are then subjected to a peak finding algorithm tailored to identifying clusters of insertions. The algorithm analyzes the frequency distribution of insertion sizes, leveraging the core idea that elements within the same TE family have similar sizes. The algorithm identifies regions with a high density of insertions as peaks of interest that deviate from the expectation, which would be SV INDELs not related to TE activity. The parameters of the algorithm are selected to balance sensitivity and specificity, although it may be difficult to fully account for variations in TE insertion densities in different species. In this case, user parameter selection is possible.

### Precursor Database Query

Based on the results of the peak finding algorithm or user input a consensus sequence is created from a subset of the insertions for each identified peak and queried against the DFAM/RepeatMasker database (Storer 2021, Smit 2010). The results of each query are output to a summary precursor file that allows the user to quickly filter for specific TE types or unannotated TEs. The user can also specify a custom set of peaks to be initially queried against DFAM (Storer 2021).

### Loci Extraction and Clustering

For each identified peak size, the corresponding sequences and loci are extracted from each sample’s VCF resulting in pseudo-merged population wide VCF for each peak size range. Importantly each sequence is retained. Each sample (within the same species) and each of its respective loci sequences are subjected to clustering analysis based on pairwise alignment sequence similarity. Each cluster of loci consists of sample specific sequences of the same length that share above the threshold of some predefined similarity score.

### Intersection with Species Genome

Each cluster for each peak obtained from the clustering analysis can be intersected with the respective species’ genome annotation file (GCF). For each intersection the gene name and intersection type (5 UTR, exon, CDS, etc.) is documented.

### Gene Annotation and Functional Enrichment

One optional step offered is to perform functional enrichment analysis on each cluster’s overlapping gene set to determine any overrepresented biological processes or pathways associated with the genes directly involved or nearby TE insertions (Kolberg 2023).

### Phylogenetic Analysis

Another optional step offered is to construct a phylogenetic tree based on one or more of the clusters. Sequences within each cluster undergo alignment, and a phylogenetic tree is constructed using maximum likelihood (Larkin 2007). These trees can elucidate the evolutionary relationships and divergence times among TE insertions within a locus. This analysis can aid in distinguishing between recently active and more ancient TE insertions especially when compared to other species.

### Output

To ensure user-friendliness and downstream analysis capabilities, two different output types are generated for each cluster and peak.

1. **Individual Information (CSV):** Each line in the CSV file contains loci information, sample name, gene intersection information, and the sequence for each member of the cluster.
2. **Merged Population Information (CSV)**: Each line in this CSV file lists individuals in the population with an insertion at a shared locus within the cluster. Instead of individual sequences, a consensus sequence for the entire cluster is established.

### Horse Case Study

As with many species closely tied to human intervention, horses present a compelling case study for TEs. Horses were domesticated in the Eurasian steppe over 4,000 years ago. The evolution of horses, owing to their pivotal role in human civilization, has left a myriad of genetic footprints influenced by various historical and cultural inflection points. As the arrival of industrialization initiated the replacement of traditional horse roles, their selective breeding was intensified, particularly for those horse breeds used for specific purposes like racing in western cultures. This has resulted in a spectrum of modern horse populations, ranging from those that have undergone extreme genetic interventions to those that experience less structured breeding and life histories. While TEs fundamental role in genome evolution is appreciated, the interplay between TEs and the genetic repercussions of diversity loss, especially in species with a varied breeding history, remains poorly understood.

To demonstrate TEPEAK, we ran it on 265 horse samples across 11 breeds, Thoroughbred, Quarter horse, Friesian, Hanoverian, Freiberger, Arabian, Mongolian, Akhal-Teke, Jeju horse, Tibetan, and Standardbred (Figure 2, Table 1), which represent a basal subset of the over 600 horse breeds. This selection was also made in part on the availability of SRA accessions on NCBI. These breeds offer diverse ancestries and histories perspectives ranging from well-documented breeding lineages to ones whose origins are subject of debate. Geographically, these breeds offer a broad representation of environmental and cultural influence. And most importantly these breeds represent a wide spectrum of human influence on horse genetics.

**Figure 2:**
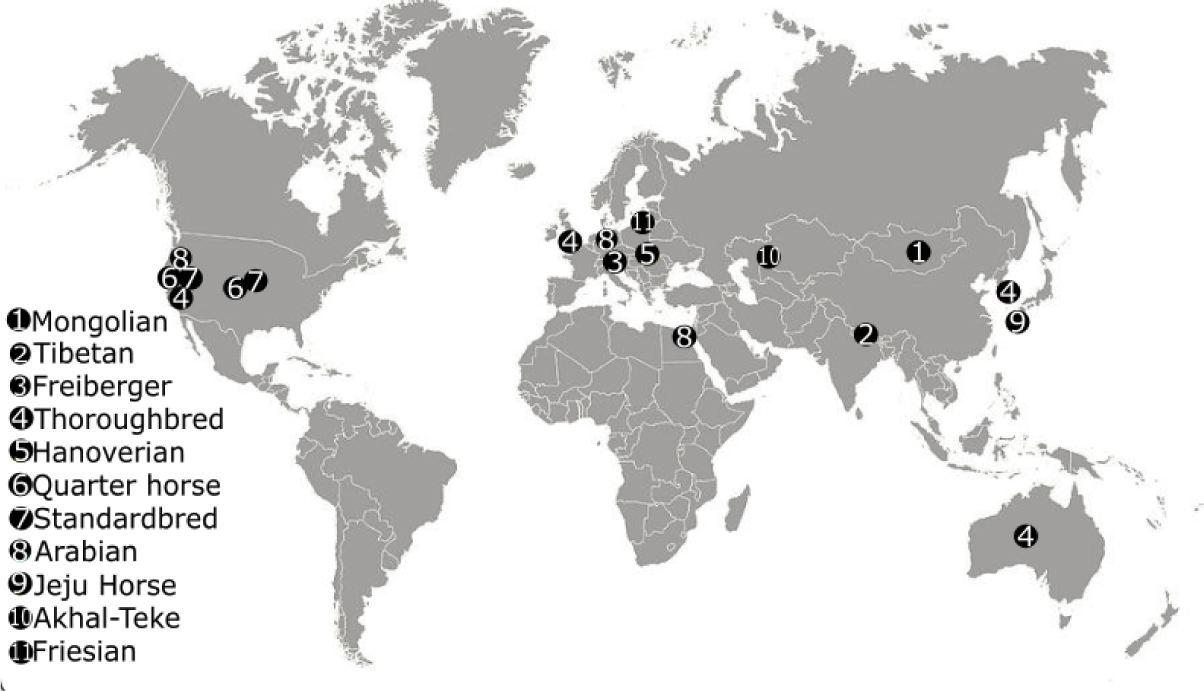
The approximate locations where each sample group was sequenced.

**Table 1:**
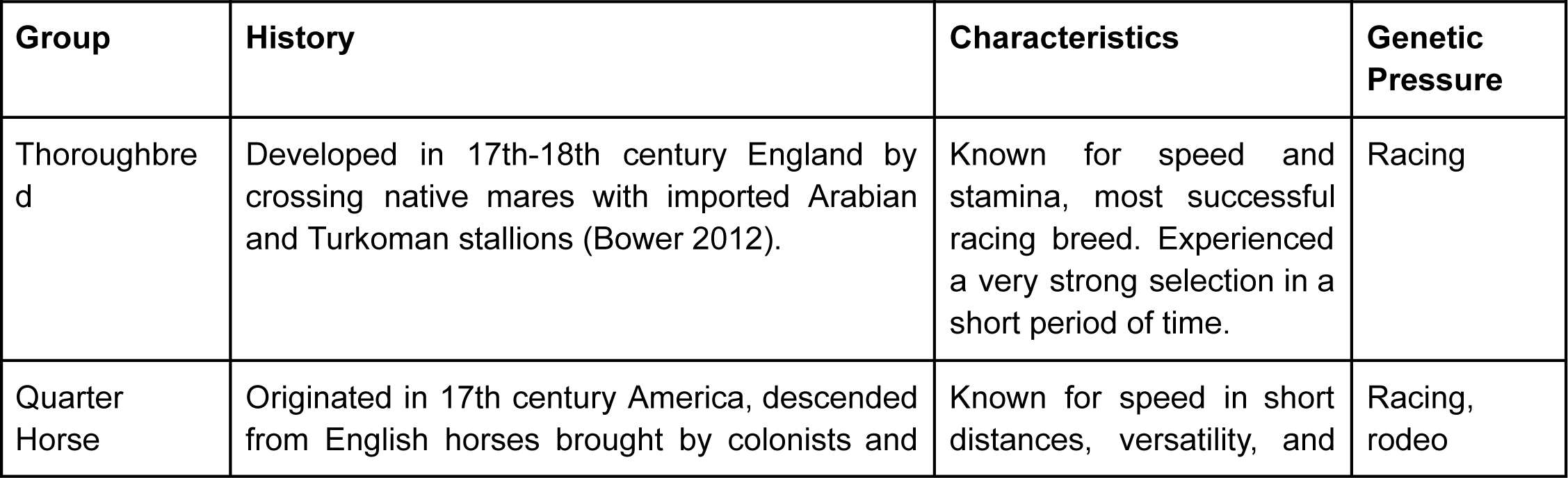
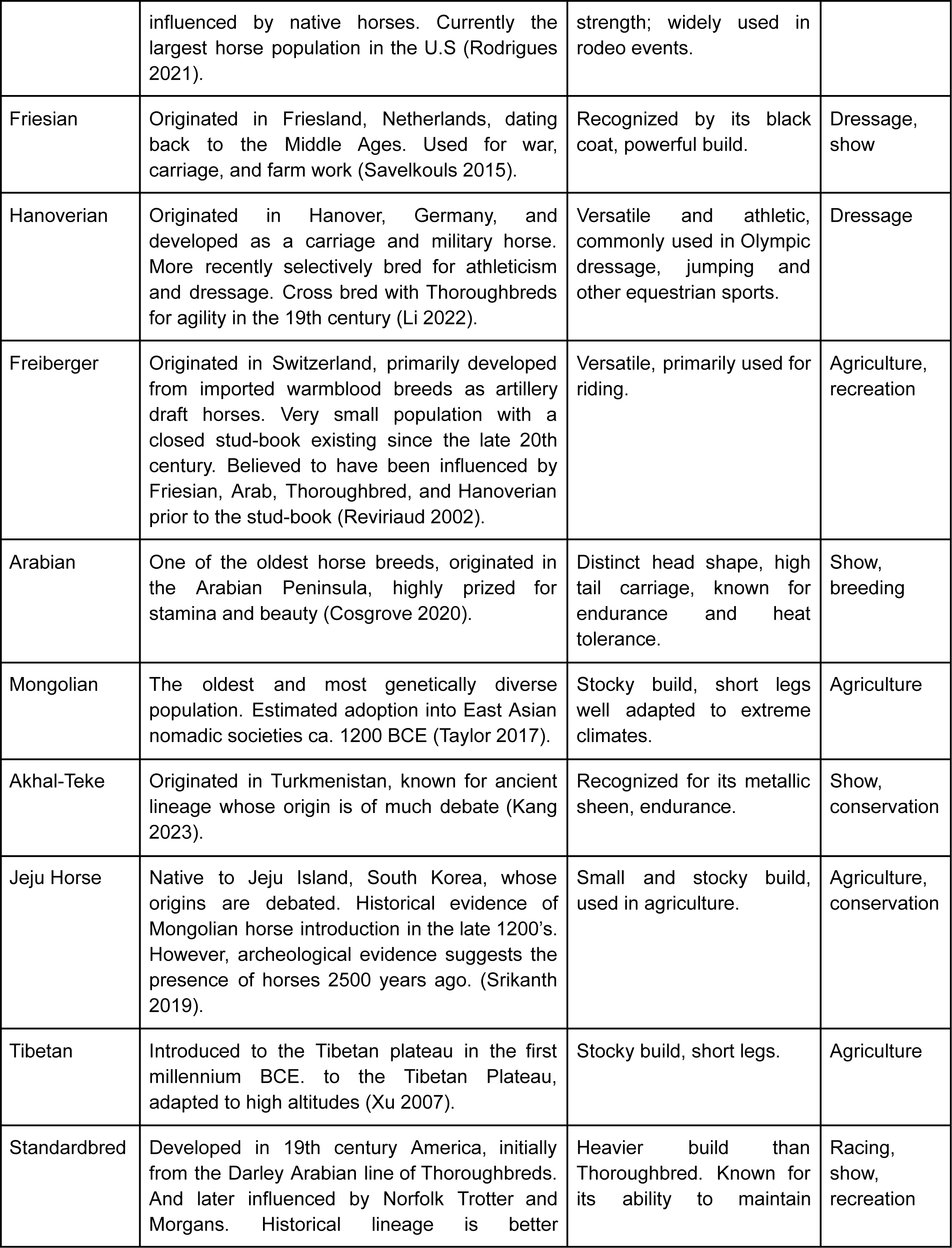
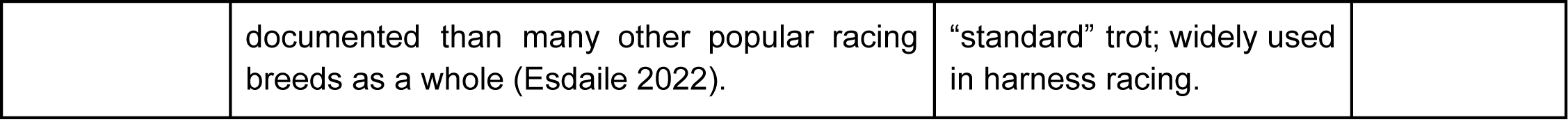
A brief history and characteristics of each group.

We selected 8 varieties of contemporary horses and three Asian landraces (Mongolia, Tibet, and Korea). Of these, several are subjected to high levels of contemporary anthropogenic selective pressure for racing and dressage (Bower 2012, Li 2022, Reviriaud 2002, Rodrigues 2021, Savelkouls 2015, Cosgrove 2020, Kang 2023, Esdaile 2022). Others, such as pastoral horses from Mongolia and Tibet, may have much lower levels of human intervention in their breeding. We expect some geographic structuring in relatedness across these varieties, with horses from Korea, Tibet, and Mongolia likely sharing an initial history of dispersal after the initial domestication of the horse in the late 3rd or early second millennium BCE (Librado 2021).

Each sample was aligned to the latest available horse genome assembly, EquCab3.0, which was isolated from a thoroughbred (Li 2009). The resulting aligned files (BAM format) were input into Insurveyor. Each sample’s VCF file was subjected to filtering to exclude insertions with a size below 100bp. Those samples with less than 1000 insertions were rejected. The resulting mean insertion count for each sample was 4312bp. The peak finding algorithm identified significant peaks at 246bp, 368bp, 490bp, 651bp, 781bp, 873bp, 1374bp, 6,400bp, 6,603bp, 6,995bp, and 7,112bp (Figure 3). To characterize the identified peaks, a consensus was established for each peak using the DFAM database. DFAM results annotated five instances as structural components of LINEs (Long Interspersed Nuclear Elements), two instances as LTRs (Long Terminal Repeats) from ERV1 (Endogenous Retrovirus 1), and four instances without existing annotations.

**Figure 3:**
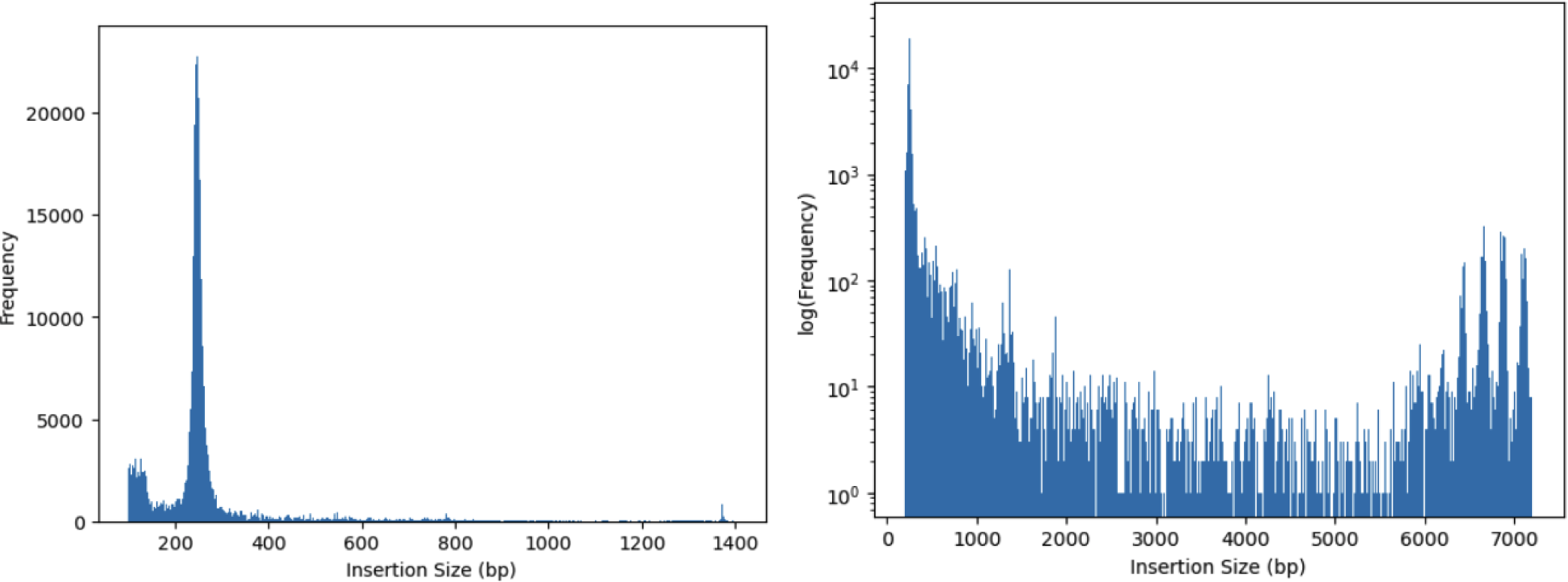
TEPEAK’s size-frequency distribution for 259 horse genomes with a linear- and log-scaled y-axis.

To further investigate peaks not annotated in DFAM, two separate approaches were undertaken. First, the sequences corresponding to these unannotated peaks were subjected to BLAST against the horse genome. In the case of the 246bp peak, BLAST identified the peak consensus against the consensus sequence for Equine Repetitive Elements 1 (ERE-1) (94% PID, 92.56% coverage). BLAST did not return any annotations for the peaks at 651bp, 873bp, and 1,374bp.

Second, to address the absence of annotations for the peaks at 651bp, 873bp, and 1,374bp, we used BLAT (BLAST-Like Alignment Tool) from the UCSC (University of California, Santa Cruz) Genome Browser. While BLAT recovered annotations for these peaks, access to many of its full annotations and consensus sequences were locked behind a subscription service, making it challenging to acquire complete information. Access to this information is not required by TEPEAK, but we aimed to utilize a validated ERV consensus sequence to show TEPEAKs ability to reconstruct full length ERVs as well as identify solo LTRs. BLAT extracted ERV1-LTR annotations for the 651bp and 873bp peaks. The full annotations and consensus sequence of the peaks were behind the paywall. The 1,374bp peak consensus sequence resulted in full coverage for the LTR regions of ERV2. ERV2-LTRs and ERV2-INT’s (internal) consensus sequence and full annotations were not behind a paywall.

ERE-1 and ERV2 were selected as the focal points for our case study. TEPEAK was run on 300 horses across 11 groups. Over 1,541,863 insertions (>100bp) passed InSurveyor’s and TEPEAK’s filtering and 35 horse samples failed to reach the minimum number of insertion calls. Insertions located on sex chromosomes were excluded.

### Extraction of ERE-1 loci

The peak finding step of TEPEAK identified a significant peak at 246bp consisting of 23,092 unique insertions. BLAST database query confirmed the insertions at this size to be a subfamily member of the perissodactyl-specific SINE family of Equine Repetitive Elements (ERE), ERE-1. To date ERE consists of four major subfamilies: ERE-1, ERE-2, ERE-3, and ERE-4 with consensus sequences ranging from 228 to 268bp (Jurka 2005).

Analysis of the size-frequency histogram showed the ERE consensus range proxies roughly 90% of prominent sizes immediately adjacent to the 246bp peak, ranging from 220-280bp. Extraction of all insertions (526,675) in the 220-280bp range and filtering of sequences using each subfamily’s consensus sequence confirmed the presence of all four ERE subfamilies. Previous work has shown ERE-1 to be the most polymorphic subfamily with strong evidence that ERE-1s were the most recently active element in the horse genome (Santagostino 2015, Gallagher 1999, Wang 2006, Sakagami 1994). Our results agree with these findings. ERE-1 insertions are not only more polymorphic in terms of interbreed and population frequency variance, but also sequence divergence and locational preference (Figure 4).

**Figure 4:**
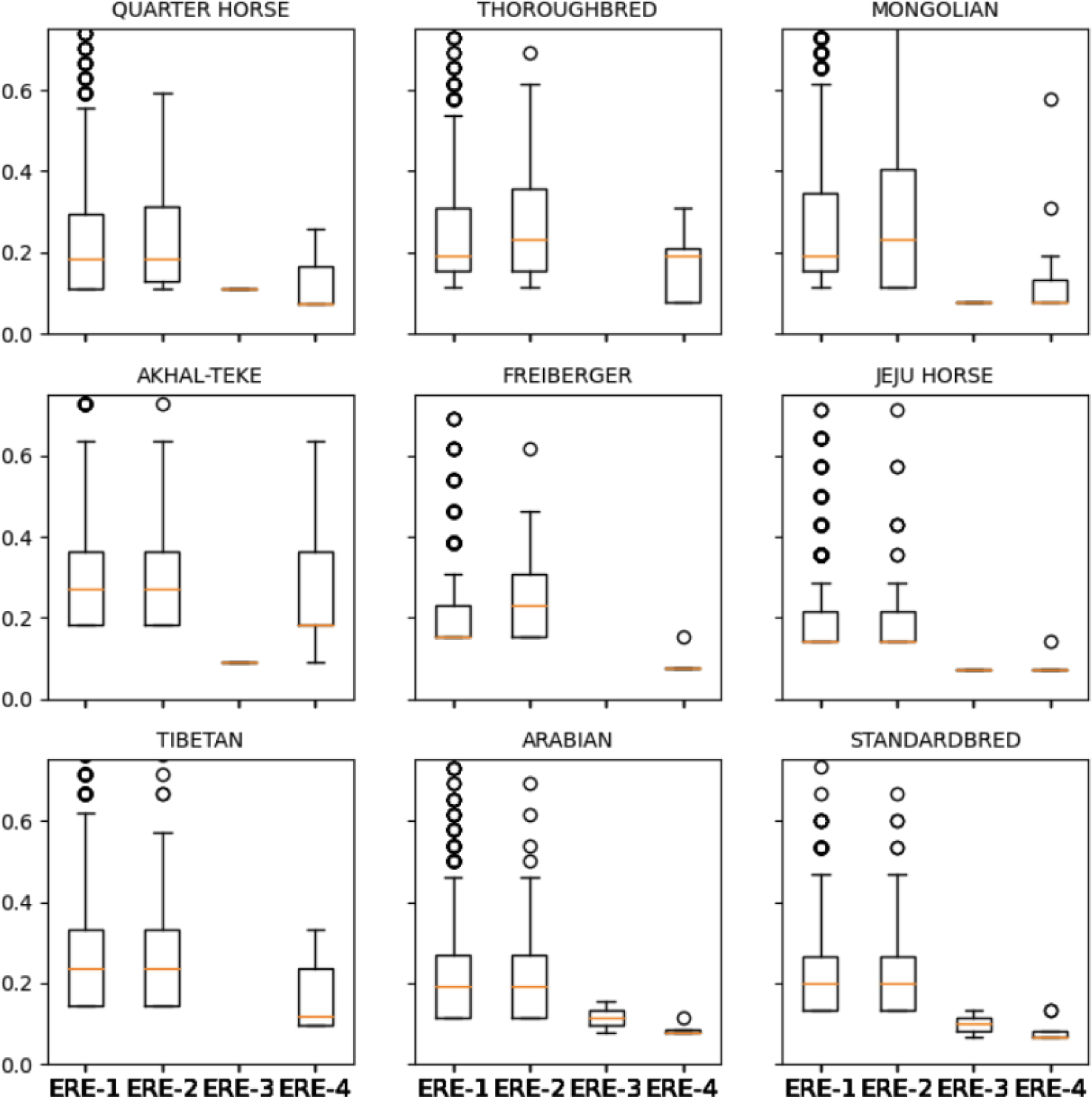
Allele frequencies for each of the ERE subfamilies with rare loci removed (less than 0.1 AF).

### Extraction of ERV2-LTR and full-length ERV2

Queries of the consensus sequence at the 1,374bp peak against the DFAM and BLAST returned no closely matching annotations. It was initially believed that this TE was a novel discovery. However, further research revealed the annotations to be owned by a subscription service, GIRI. Utilizing UCSC BLAT, which has access to the GIRI RepBase, to query the 1,374bp consensus sequence against the horse reference confirmed this TE to be the long-terminal repeat region of an endogenous retrovirus, specifically ERV2-1N-EC_LTR (ERV2-LTR) (Jurka 2005). ERV’s contained open reading frames (three genes) when initially infecting hosts millions of years ago. This internal region is flanked by identical long terminal repeats (LTRs). Over time the internal region becomes inert and often becomes excised through recombination leaving the two LTRs in close proximity to each other (Wells 2020, Yu 2013). It is estimated that 90% of all ERVs are solo LTRs (Chuong 2017, Wells 2020).

We then demonstrated TEPEAK’s ability to extract full-length and intact ERVs without prior knowledge of the internal region sequence. While a full-length ERV may be observed as a significant peak in the size-frequency histogram, the polymorphic nature and similar size to much more common LINE insertions make it difficult for de-novo extraction as was performed on ERE-1. To demonstrate this we first extracted and filtered all ERV2-LTR loci by pairwise alignment with a consensus sequence constructed from all insertions in the 1,370-1,380bp range. Any sequence with less than 85% PID was rejected. This filtering extracted 2,483 insertions over 544 loci. Subsequently, we merged all loci within a distance of 10,000bp to one another. This threshold was a deliberate overestimation for the interior region length in order to account for instances where paired LTRs might not have been successfully identified as well as mitigating any other resolution inconsistencies in repetitive regions. This merging filter reduced the number of loci to 200. It is worth noting the average distance of the original loci prior to merging was less than 50bp. The merged loci were then intersected against the entire population’s insertions whose size was greater than 4,000bp. This threshold is a conservative underestimate for the internal region of the ERV. Finally, the resulting sequences were split in half where each half was scanned against the ERV2-LTR consensus sequence. A successful match in both halves was classified as full length ERV.

## Results

### Polymorphic insertions of ERE-1 shows distinct breed population structure

As ERE-1 represents the most recently active ERE element we chose to focus on just ERE-1’s population structure. We thus isolated insertions with sequences that share at least 85% identity to the ERE-1 consensus. This filtering extracted 22,451 loci and 285,828 insertions. Across all breeds, a majority of ERE-1 loci are rare (less than 0.1 AF), 2,900 loci were in at least 50% of the population in any breed, and 116 loci were in at least 50% of all breeds. UMAP results validate our intuitions for most breeds (Figure 5B). Notably, the Jeju horse, isolated on an island and not subjected to racing or dressage breeding pressure, serves as a pseudo control group.

**Figure 5:**
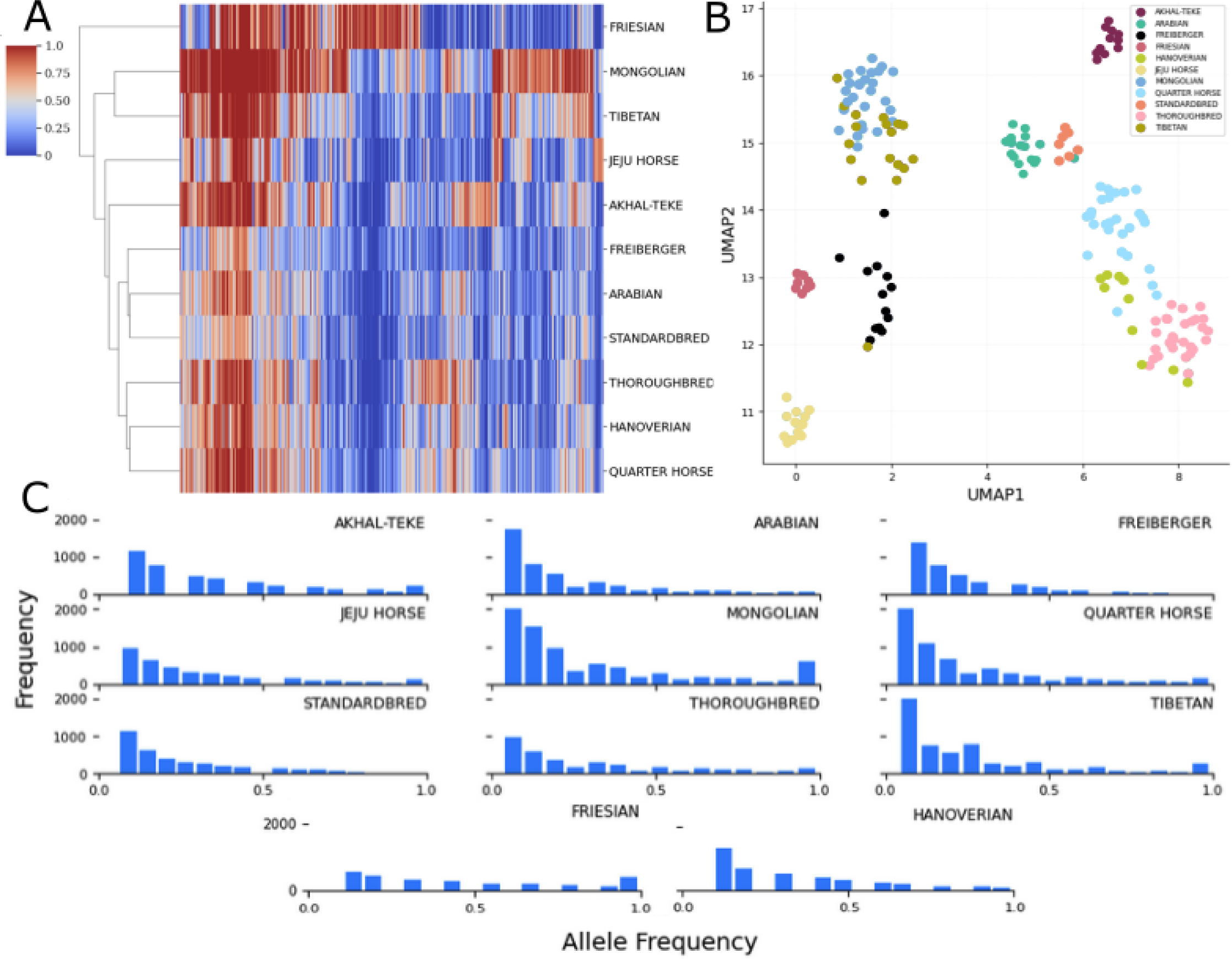
A. A heatmap of the allele frequencies of individual ERE-1 insertion loci (x-axis) across the horse breeds (y-axis). Only loci with a minimum allele frequency of 0.75 in at least one of the horse breeds are included. Color scale indicates allele frequency in each breed, the transition from blue to red indicates ERE-1 insertions in the majority of samples for the respective breed. The y-axis represents the resulting dendrogram from hierarchical clustering. B. Visualization of UMAP dimension reduction on allele frequency data based on alleles that are present in at least 0.75 of one horse breed. C. Inter-breed allele frequency distribution for all ERE-1 loci present in at least one sample belonging to the respective breed.

Thoroughbred and Arabian breeds exerted a discernible influence on the genomic makeup of Quarter Horse, Standardbred and Hanoverian, as evidenced by their grouping patterns. However, a distinct dissimilarity was observed in the Freiberger population despite the evidence of some shared Thoroughbred lineage. Surprisingly, despite the influential role of Thoroughbred and Arabian breeds, both exhibited relatively few nearly fixed (> 0.75 AF) ERE-1 loci, appearing to have passed on only a limited subset of nearly fixed loci. To further understand this subset of shared loci, we filtered all loci shared by at least 75% of individuals within each breed (Thoroughbred, Arabian, Standardbred, Quarter Horse) and annotated their sites for genomic features. The initial filtering extracted 88 loci, 27 of which intersected with genomic features (10 intronic regions, 4 exon regions, 12 unknown horse genes, and one enhancer region). [Table 2]. Curiously, 25 of these loci are also nearly fixed in the Mongolian population. The intersected genes highlight critical developmental, behavior, and physiological processes associated with memory, temperament, neurological and vascular systems, as well as major biological metabolic and synthetic pathways. We acknowledge insertions into introns may be nonfunctional unless demonstrated otherwise experimentally. However, variation in heritability of these shared loci especially in breeds with drastic selective breeding pressures is notable due to the greater chance of viable and stable insertions (Kidwell 1997). This notion also demonstrates TEPEAK’s ability to filter for genomic targets in additional analysis.

**Table 2:** Genes associated with the nearly fixed loci shared among Mongolian, Thoroughbred, Standardbred, Quarter Horse, and Arabian breeds. All related traits sourced from GeneCards unless specified otherwise.

| Gene Name | Related Traits | Insertion Type |
| --- | --- | --- |
| ALDH18A1 | Temperament (Benítez-Burraco 2020) | Intron |
| KIF11 | Spindle Dynamics/Dendrites<br>Overexpression believed to offset<br>Alzheimers in humans (Lucero 2022) | Exon |
| ALK | Insulin receptor/brain development | Exon |
| SYN3 | Synaptogenesis/modulation of<br>neurotransmitters | Intron |
| Enhancer for MEIS1 | Hox Gene/head-tail axis development | N/A |
| ZNF121 | Predicted DNA binding TF activity | Intron |
| Unknown Horse Gene<br>ENSECAG00000057187 | UNK | UNK |
| MyH10 | Cell maintenance with roles in coronary, vasculature function (Kim 2018) | Intron |
| GUCY1B2 | GTP and heme binding | Exon |
| Rag1 | Immunoglobulin V-D-J recombination | Intron |
| AUTS2 | Neurodevelopment, candidate gene for numerous neurological disorders | Intron |
| DCLK1 | Memory and general cognitive abilities | Intron |
| LRP1B | Clearance of extracellular low density lipids | Intron |
| PALLD | Actin cytoskeleton | Intron |
| IGF2BP3 | RNA synthesis - late development | Intron |

There is a direct correlation between breeds subjected to selective breeding pressure and the prevalence of both rare and fixed loci. This relationship highlights the genetic loss that results from such practices. Intriguingly, the Quarter Horse appears to deviate from this trend. This possibly could be attributed to the high introgression of other lineages as well as its extensive population in the United States. The Quarter horse’s connection to Spanish, Native American, and Anglo breeds, many of which are not explored in this study, suggests unique genetic dynamics from the different populations and encourages a future exploration..

### Mongolian horses as a source of ERE-1 alleles

The Mongolian breed is the oldest and most genetically diverse contemporary horse population (Kusliy 2021). It is thought that a majority of SINEs amplified at least 30 million years ago in proto-ancestors of mammals (Marchani 2009, Etchegaray 2021). Although ancient DNA suggests that horses were first domesticated in western Eurasia, early horses dispersed to the region as early as ca. 1200 BCE or before (Taylor 2017) and Mongolia likely played a key role in the initial dissemination of horses into nearby regions of East Asia (Rawson 2021). Beginning in the first millennium BCE and continuing through the Middle Ages, expansive pastoral empires like those of the Xiongnu, Turkic Khaganate, and the Mongol Empire expanded the role of Mongolian horses across much of Eurasia.

The presence of most of the variation in the Mongolian population implies that the ERE-1 loci were extant at the time of breed diversification. It is also apparent that this variation continues to exert an influence on modern-day horse genetics despite strong selective breeding in many populations.

The Mongolian breed exhibits the greatest amount of fixed and rare ERE-1 loci, highlighting the intuition that the Mongolian breed is the best representation of ERE-1’s impact on an intact population. To further investigate the shared relationship between Mongolian and other modern-day breeds, we assessed the number of shared loci in pairs of horse breeds, considering instances where at least 75% of samples in each breed exhibited insertions at each loci (Figure 6). These results affirm the Mongolian breed as harboring the most diverse set of loci. Furthermore, we observed a notable pattern in the distribution of shared loci, particularly among breeds in proximity to the Eurasian steppes.

**Figure 6:**
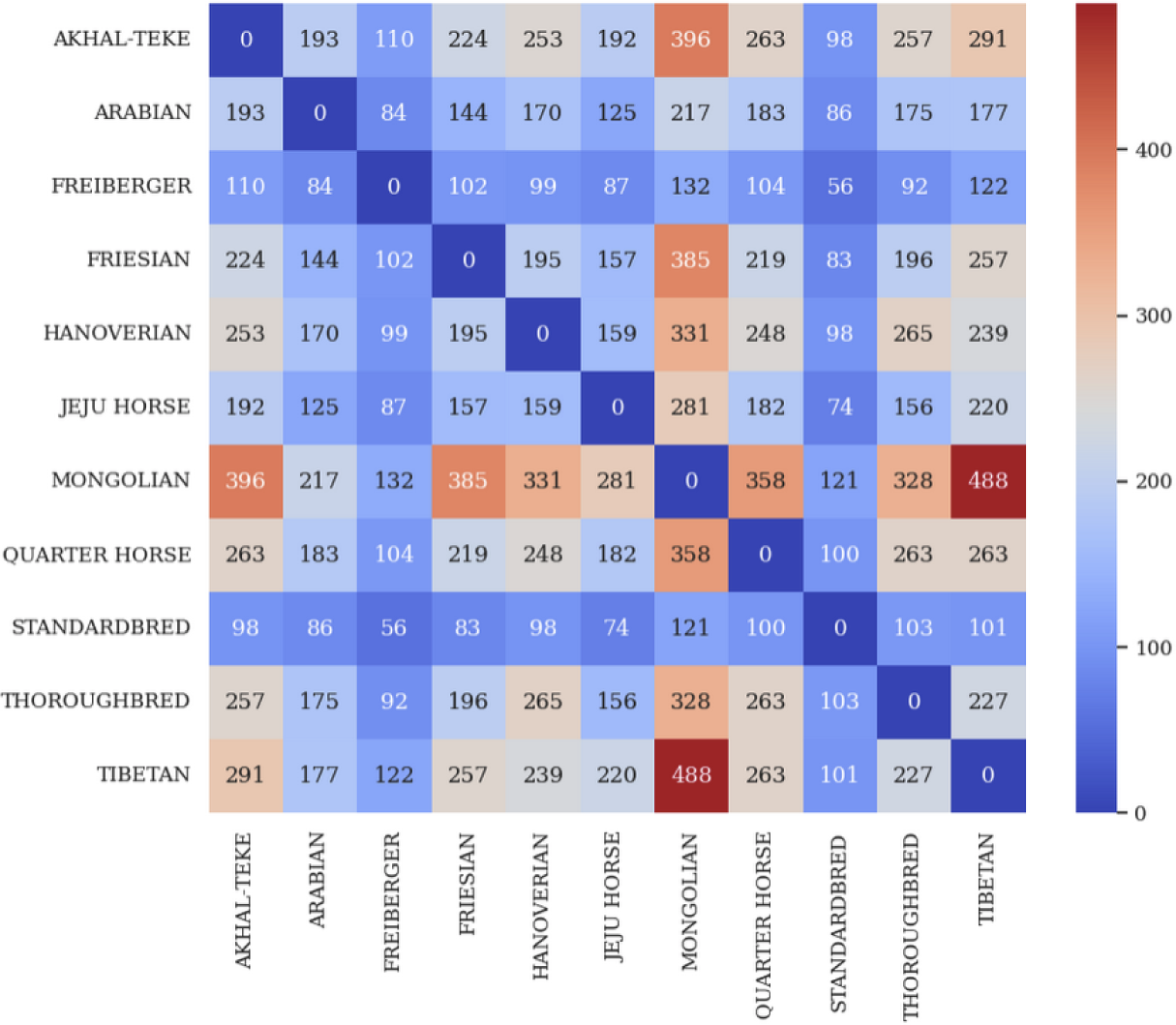
The number of pair-wise shared high-frequency (AF>=0.75 in both groups) loci.

A considerable amount of shared loci between Mongolian and Akhal-Teke was also observed. These results point to shared history between the two varieties, and encourage further exploration into ancient connectivity across Inner Asia incorporating ancient genomes.

As discussed in the preceding section, we emphasized the limited number of nearly fixed loci shared among racing and Mongolian horses, which are closely tied to crucial genes. Antithetical to this we observed a number of loci that are variable in Mongolian populations, but have become fixed in selective breeds. Here we explore the number of nearly fixed loci shared among racing breeds, but not Mongolian. This filtering extracted 2 intron and 1 exon insertions into different genes (Table 3). The intron loci relate to bone and growth development regulation and the exon antigen production. The loci that are polymorphic Mongolian and Tibetan breeds and fixed loci in nearly all race breeds could be explained by a founder effect and selective breeding.

**Table 3:** Genes associated with the polymorphic loci in Mongolian and Tibetan, but fixed in Thoroughbred, Standardbred, Quarter Horse, and Arabian.

| Gene | Mongolian | Tibetan | Akhal-Teke | Thoroughbred | Standardbred | Quarter Horse | Arabian |
| --- | --- | --- | --- | --- | --- | --- | --- |
| BBX | 0.38 | 0.28 | 1.0 | 1.0 | 0.8 | 0.85 | 0.84 |
| BTN1A1 | 0.42 | 0.43 | 0.36 | 1.0 | 0.86 | 0.85 | 0.76 |
| PTPN2 | 0.42 | 0.28 | 0.82 | 1.0 | 0.8 | 0.81 | 0.76 |

### Genomic Impact/Functional Enrichment of ERE-1

Next, in order to highlight TEPEAK’s ability to gauge the overall genomic influence of a TE, we performed an annotation and functional enrichment analysis of all ERE-1 loci. The results of this intersection pinpointed the presence of ERE-1 in 56 coding DNA sequences (CDS), 5,785 intronic regions, and 539 5’ or 3’ UTRs. 55.3% of these genes have been subject to two different ERE-1 loci. Of the CDS loci several significant genes were noted including collagen production and joint health (COL1A2), skeletal muscle development (WASHC5), significant regulator of Hox expression-vertebrate trunk development (Nr6a1), anti-bacterial defense (C8A), smooth muscle vasoconstriction (EDNRA), neuroprotein degradation (UBQLN1), epithelial sodium channel regulation (CNKSR3), and various DNA and RNA processing and regulation (BTG4, ATG14, WDR75, TBCD, DTWD2, DDX31).

We subsequently performed functional enrichment analysis on all the CDS and exon intersected genes. ERE-1’s presence is notably pronounced in several cellular and biological pathways (Kolberg 2023) (Figure 7).

**Figure 7:**
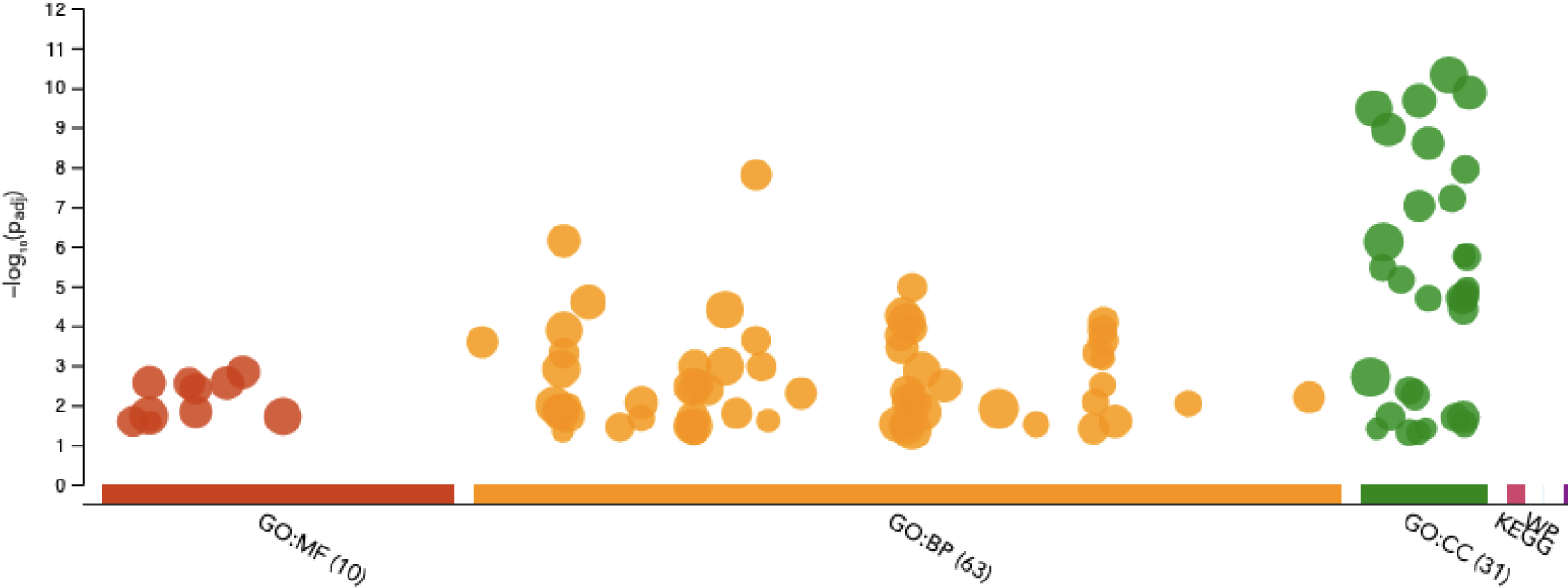
Functional enrichment analysis of exon and CDS intersected genes across all eleven horse breeds. Y-axis displays adjusted p-value, x-axis categories segregates Gene Ontology (GO) categories: Molecular Function (MF), Biological Process (BP), and Cellular Component (CC).

We also aimed to explore the potential for evolutionarily conserved relationships of these functional loci across three breeds: Mongolian, Quarter Horse, and Thoroughbred (Figure 8), chosen because of their large sample size, distinct UMAP clusters, and varying degrees of perceived selective breeding influence. Preliminary observations suggest that, despite intronic regions being conserved, the Mongolian loci experienced much higher rates of fixed intronic loci (>200) compared to the other breeds which maintained more comparable rates of insertion. Quarter horses had the highest amount of rare and common (0.1-0.5 AF) intronic loci, exceeding Thoroughbred by over 250 loci. These results validate our understanding that intronic insertions are most often highly conserved.

**Figure 8:**
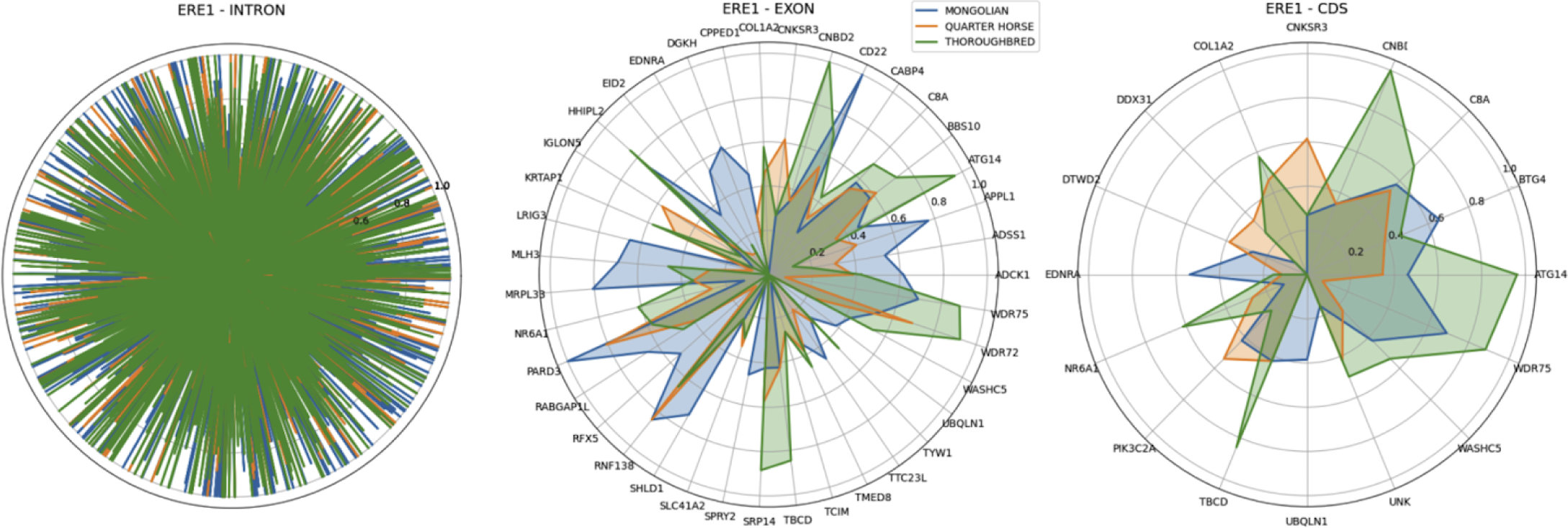
ERE-1 Allele frequency dynamics across different genomic regions, intron (left), exon (middle), and CDS (right), of three horse breeds. X-axis displays human gene orthologs and y-axis breed-specific allele frequency. Intron gene orthologs are omitted for clarity.

In exonic loci both the Thoroughbred and Quarter Horse displayed matched patterns of insertions, with the Thoroughbred showing higher rates across most genes. The Mongolian horse was observed to have smaller rates of exonic insertions compared to the intronic loci, however displayed a more unique set of loci that appear to be rare or polymorphic in the other breeds. This is undoubtedly an artifact of the Mongolian horse’s maintained heterozygosity. In the CDS loci the Thoroughbred was observed to have higher rates of fixed and nearly fixed loci deviating from both Mongolian and Quarter horse in many genes, suggesting another consequence of reduced genetic variation from selective breeding practices.

### Extraction of full-length endogenous retroviruses in horse

TEPEAK was able to extract 31 different full-length ERVs across 10 loci. Multiple sequence alignment of each ERV against RepBase’s consensus sequence for ERV2_INT (4,450bp) validated the presence of the intact internal region in all 31 insertions with no false positives. In order to validate the sensitivity of this method we performed pairwise alignment of ERV2_INT’s consensus sequence against all insertions over 4,000bp (75,623 sequences) (Jurka 2005, Larkin 2007). This brute-force method is computationally expensive. This method yielded 5 additional full length ERV sequences that matched our previously discovered 10 loci. It also yielded three new loci with three insertions total. Further analysis showed two of the new loci to have valid full-length ERVs. However the third loci, which only contained one full length insertion, contained one LTR region whose sequence shared poor identity to the ERV2-LTR. Additionally, this loci matched none of the filtered loci extracted from the 1,370bp-1,380bp range. Therefore this loci was rejected as not a full length ERV2. Furthermore it was observed that the two new validated loci contained shorter LTR regions (1,290bp-1,340bp), which consequently resulted in rejection from the previous method.

We also annotated each of the ERV2 (both LTR and full length) loci for genomic features, resulting in intersections with 112 different genes, 1 CDS, 14 UTR, and 97 intron. It was observed that despite intersecting with critical genes (RPH3A (intron), GABRR1 (5’UTR), SVC2 (intron), STXBP5L (intron), and CDK20 (5’UTR)), the loci where full length ERV2 was identified made up a set of the least conserved loci across the species (Figure 9). The loci that make up the most conserved loci across the species intersect with (AVEN (intron), Igλ (intron), HLA (5’UTR), MHC/EQMCE1 (intron), and ZNF77 (5’ UTR)). We expect to observe immune system related genes to be enriched with TE activity, however it is notable that the nearly fixed loci for these genes are most present in Eurasian horse breeds (Jeju horse, Akhal-Teke, Mongolian, Tibetan, Arabian).

**Figure 9:**
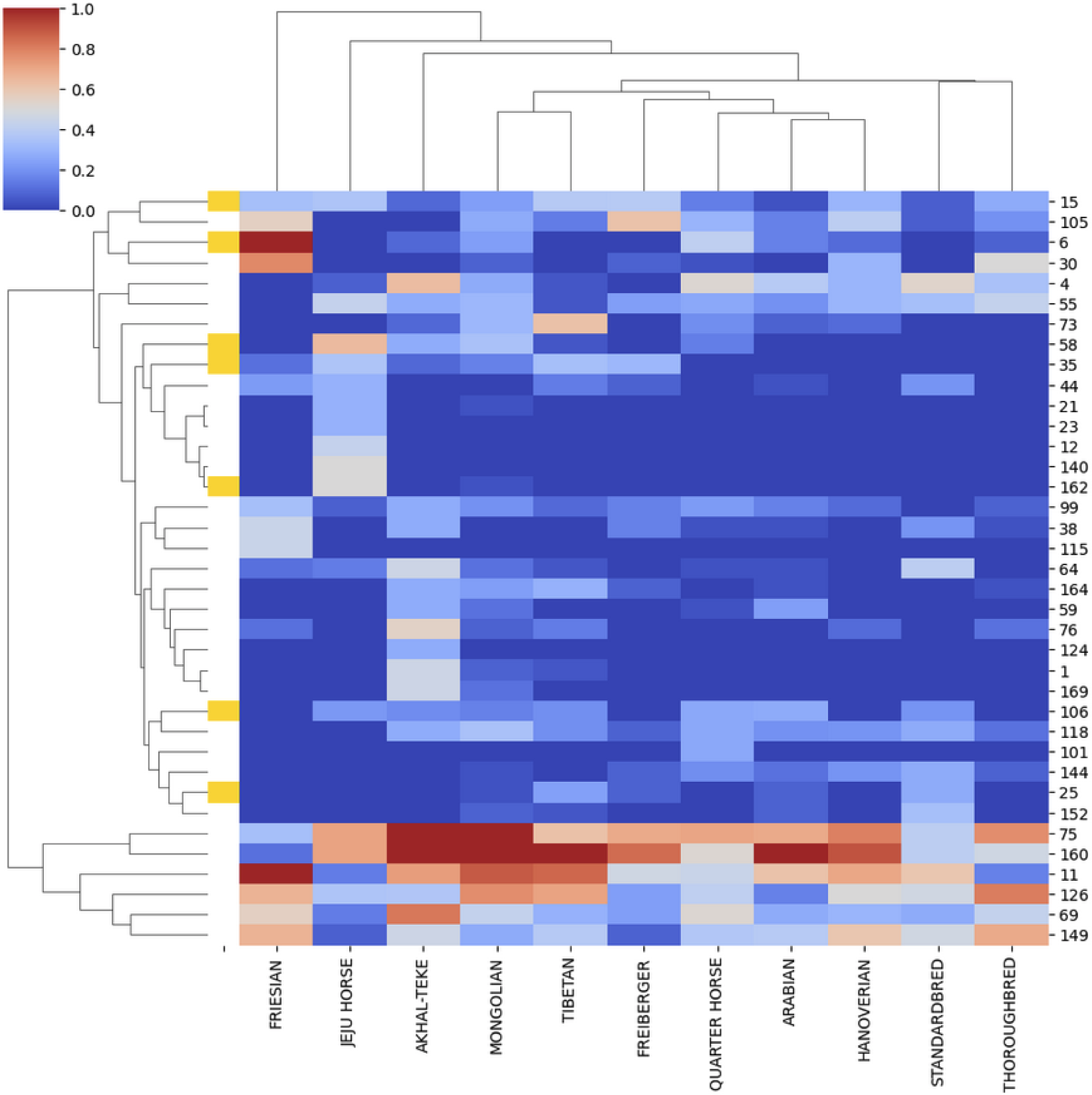
ERV2-LTR loci’s allele frequency in each breed. Loci that are not present in at least 40% of one breed are excluded. The orange bars on the left mark loci where full length ERV2s were found. Notable gene intersections with loci enriched in the population: 75: EQMCE1/MHC (intron), 160: Igλ (intron), 11: AVEN (intron), 126: unknown horse gene, 69: HLA (exon), 149: ZNF77 (exon).

We then filtered the sequences from each of these four species within the 220bp-280bp range and constructed consensus sequences. Pairwise alignment of each species’ consensus with the horse ERE-1 consensus resulted in >99% for zebra and wild ass. However black rhino and Brazilian tapir were observed to have 76% and 74% PID. We then compared the horse ERE-2, ERE-3, and ERE-4 consensus sequences to the rhino and tapir. In both species the ERE-3/4 resulted in the highest identity, >94%. This further validates the evidence that ERE-1 is the oldest member of the ERE family. We were unable to identify any insertions with high identity (>75% PID) to ERE-1 in rhino or tapir. To our knowledge, wild ass is the only species tested with annotations for this particular sequence, labeled as ERE-1. [Table 4]

**Table 4:**
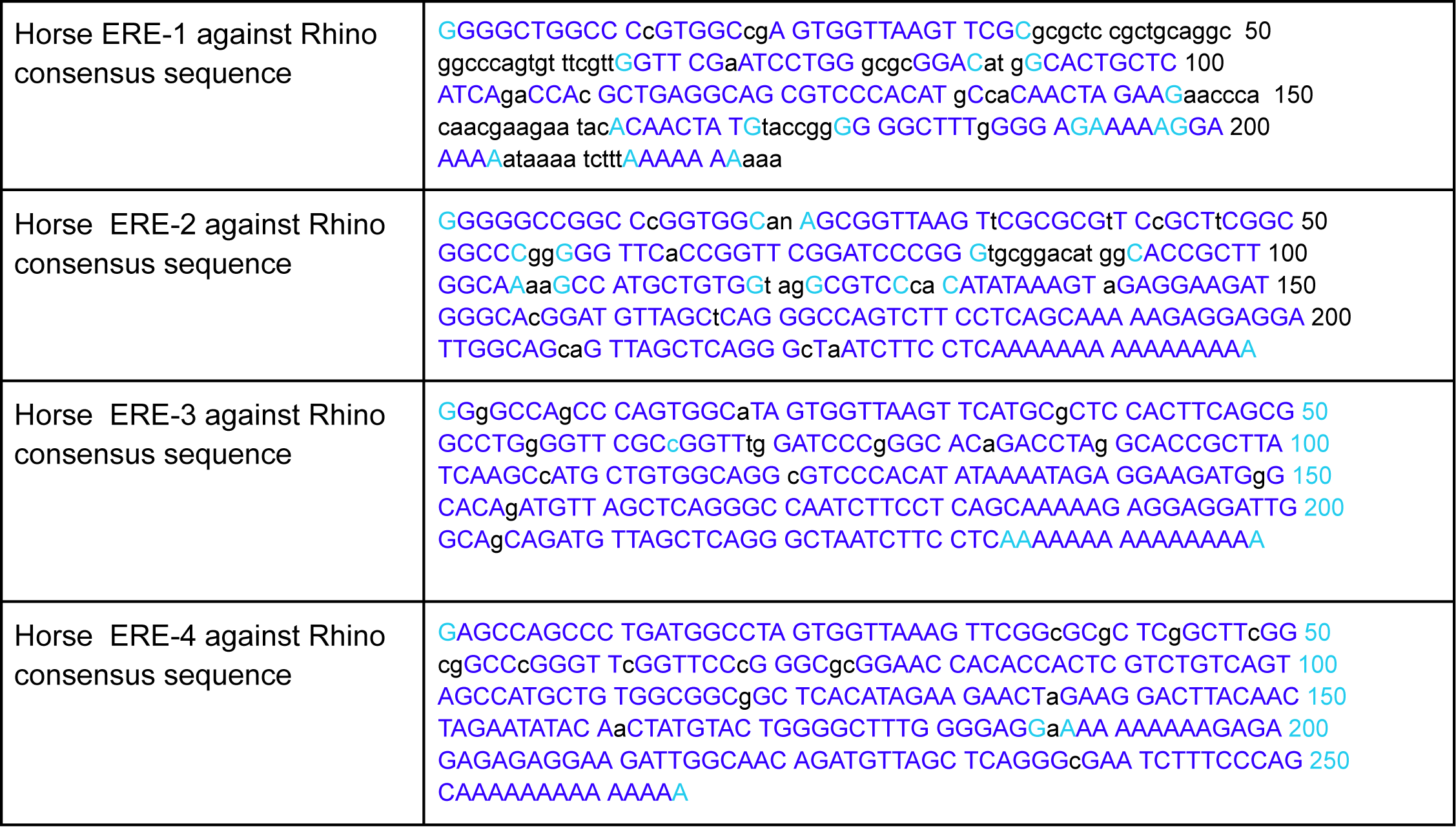
Pairwise sequence alignment results of each of the horse ERE consensus sequences against the Rhino 220bp-280bp consensus sequence. Capital letters indicate matches and lowercase letters indicate mismatches. Light blue represents GAP boundaries.

Despite the limited insertions in the range of ERV2-LTR, we were able to identify sequences with high PID (>98%) in both wild ass and zebra, but not rhino or tapir. Additionally, due to the low quality reads in these samples no fully-intact ERV2 were able to be extracted. However, BLAT of the horse full ERV2 sequence resulted in sequences with high PID and query coverage, 95% and 92%, in both wild ass and zebra. Neither the LTR nor the full ERV regions are annotated in any database to our knowledge (Table 5).

**Table 5:** Summary of shared equid TE families found in the odd-toed ungulate species and the status of their annotations.

|  | ERV2-LTR |  | ERE-1-4 |  |
| --- | --- | --- | --- | --- |
| Species | Present? | Annotated? | Present? | Annotated? |
| Horse | Yes | Yes | Yes, ERE-1-4 | Yes |
| Zebra | Yes | No | Yes, ERE-1-4 | No |
| Wild Ass | Yes | No | Yes, ERE-1-4 | Yes |
| Black Rhino | NO | N/A | Yes, ERE-2-4 | No |
| Tapir | NO | N/A | Yes, ERE-2-4 | No |

### Novel TE in Black Rhino

Based on the analysis above, we believe that we have identified a novel TE family in black rhino of size 535/536bp. Due to the proximal nature of insertions in this range, it was hypothesized that these insertions to be ERV-LTRs. With this intuition TEPEAK was able to extract two loci with possible full length ERVs (6705bp). BLAST results of this sequence showed it to be a Gammaretrovirus (98% coverage, 92.8% PID), validating this TE family to be an ERV.

## Discussion

TEPEAK is a framework designed to identify and characterize TEs in populations using concentrations of similarly sized insertions (Figure 3). TEPEAK’s strengths lie in its ability to broaden the analysis scope of a project by integrating previously sequenced SRA samples, discovering TEs without prior annotation or sequence knowledge, and aiding users in discerning the population structure and functional impact of the TEs they find. With these capabilities, we expect TEPEAK to be particularly useful for wild populations where much less is known about TE activity and function.

To demonstrate TEPEAK and to serve as a guide for other TE explorations, we presented an analysis of TEs in 265 horses across 11 different groups and breeds, which included the related species zebra, wild ass, black rhino, and tapir. Separately, we also ran TEPEAK on giraffes, hippos, panthers, camels, cows, sperm whales, and gray whales (Supplementary Figures X-Y, Supplementary Figures I-J). TEPEAK’s SRA interface makes this exploratory, population-scale analysis feasible and efficient, which is critical for fully understanding novel TEs in unstudied populations.

Among the TEs we found in horses, ERE-1, was observed to have a set of fixed loci shared by all groups despite complex history. Despite this shared set, we also identified a number of insertion sites into the coding regions of genes coding for traits such as joint health, skeletal muscle development, and heart function. These results suggest that ERE-1 plays an active role in modern day horse genetics and breeding. We also found ERV2 insertions in both LTR and full-length form that affected immune genes and were nearly fixed in Eurasian horse breeds (Jeju horse, Akhal-Teke, Mongolian, Tibetan, and Arabian).

While TEPEAK is easy to use and powerful, it does require high-quality samples to run effectively, which can be challenging when considering ancient genomes. We also only consider Illumina short-read data sets. As long-read sequencing continues to become more popular, this could limit TEPEAK’s utility. Our automated identification and characterization methods can also struggle to correctly separate overlapping families in the same size region and may require some manual curation (e.g., removing LINEs from LTRs).

## Supplementary Tables

Supplementary Table 1: All Horse SRA Samples

https://drive.google.com/file/d/1YY-V6zhLOTQsOdQSn2Axqnv-SRHu6i5E/view?usp=sharing

Supplementary Table 2: All ERE1 Genes

Intron: https://drive.google.com/file/d/1pMiO8zA1ozLdtHt3vFhMErNUJ2GHpjL-/view?usp=drive_link

Exon: https://drive.google.com/file/d/15sHVmbFVZCvog7KoO6r5ZD9Q6dTYM-nO/view?usp=drive_link

CDS: https://drive.google.com/file/d/1W74ki8MeVqMbnhXvT3f857wC-cpv2Dsu/view?usp=drive_link

Supplementary Table 3: All LTR Genes

Intron: https://drive.google.com/file/d/1ZNPsRqxHkXkRwjp3G3fMTTmcJZRUM7sm/view?usp=drive_link

Exon: https://drive.google.com/file/d/1Bj2bI0LDxWCcbJjac_a4jUAtNSj6zffP/view?usp=drive_link

CDS: https://drive.google.com/file/d/14CBdbxgjLsSd-10qx-vhYmohgcMwMXpu/view?usp=drive_link

## Supplementary Figures

**Supplementary Fig 1:**
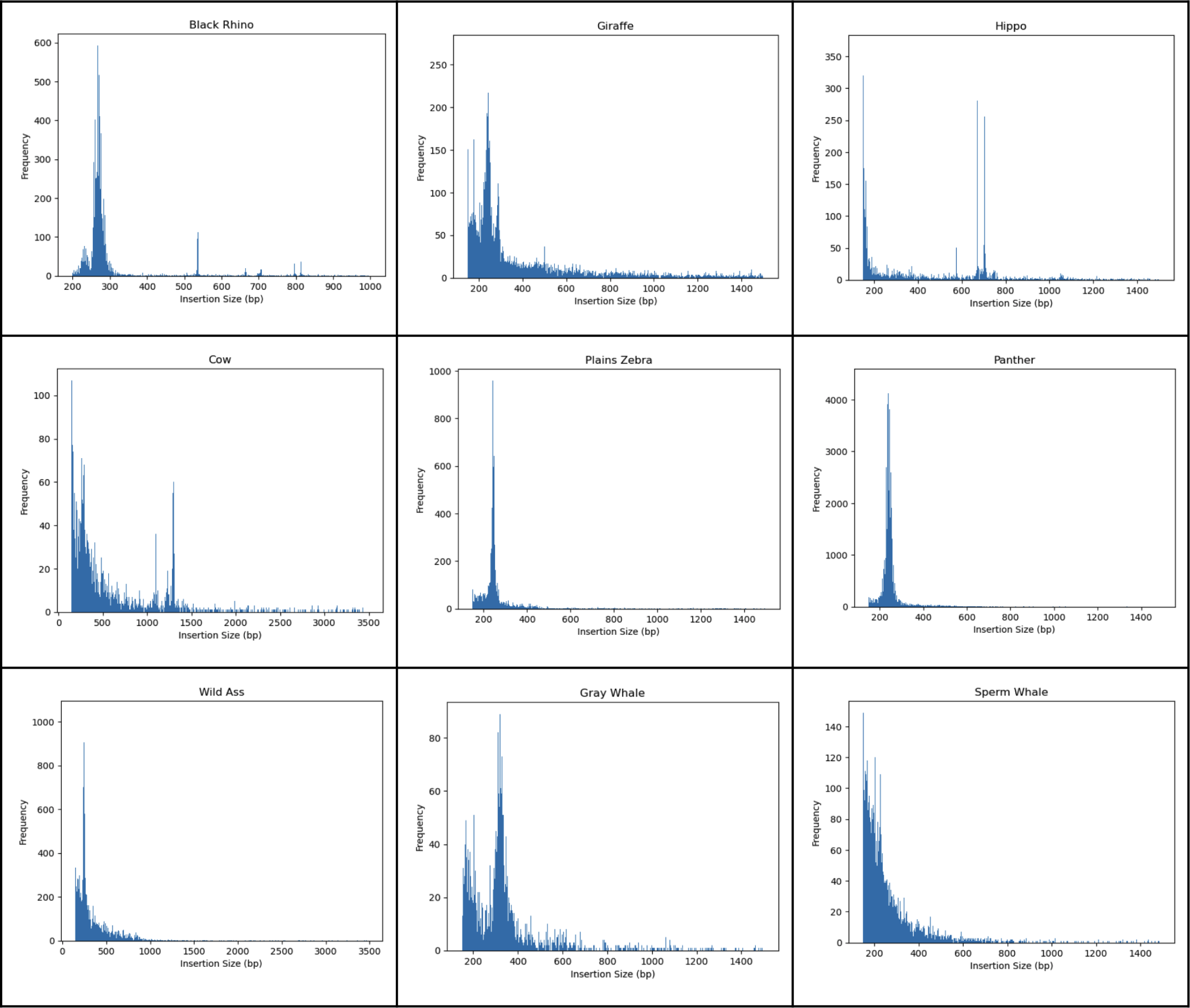
Other species size frequency distributions.

